# Potassium effects on NCC are attenuated during inhibition of Cullin E3-ubiquitin ligases

**DOI:** 10.1101/2021.11.30.470531

**Authors:** Sathish K Murali, Robert Little, Søren B Poulsen, Mohammed Z Ferdaus, David H Ellison, James A McCormick, Robert A Fenton

## Abstract

The thiazide sensitive sodium-chloride co-transporter (NCC) plays a vital role in maintaining sodium (Na^+^) and potassium (K^+^) homeostasis. NCC activity is modulated by the with-no-lysine kinases 1 and 4 (WNK1 and WNK4), the abundance of which are controlled by the RING-type E3 ligase Cullin 3 (Cul3) and its substrate adapter Kelch-like protein 3. Dietary K^+^ intake has an inverse correlation with NCC activity, but the mechanism underlying this phenomenon remains to be fully elucidated. Here, we investigated the involvement of other members of the Cullin family in mediating K^+^ effects on NCC phosphorylation (active form) and abundance. In kidneys from mice fed diets varying in K^+^ content, there were negative correlations between NCC (phosphorylated and total) and active (neddylated) forms of Cullins (Cul1, 3, 4 and 5). High dietary K^+^ effects on phosphorylated NCC were attenuated in Cul3 mutant mice (CUL3-Het/Δ9). Short-term (30 min) and long-term (24 h) alterations in the extracellular K^+^ concentration did not affect Cullin neddylation levels in *ex vivo* renal tubules. Short-term, the ability of high extracellular K^+^ to decrease NCC phosphorylation was preserved in the presence of MLN4924 (pan Cullin inhibitor), but the response to low extracellular K^+^ was absent. Long-term, MLN4924 attenuated the effects of high extracellular K^+^ on NCC phosphorylation and responses to low extracellular K^+^ were absent. Our data suggest that in addition to Cul3, other Cullins are involved in mediating the effects of K^+^ on NCC phosphorylation and abundance.

## Introduction

Hypertension or high blood pressure (BP), affects more than 20% of the population and is a major public health problem due to its contribution to stroke, heart failure and kidney failure [1]. Although sodium (Na^+^) rich diets are major contributors to the onset of hypertension, a low potassium (K^+^) intake often accompanies this high Na^+^ intake, and dietary K^+^ intake can have an inverse correlation with BP [2, 3]. In the kidney, the thiazide-sensitive sodium chloride cotransporter (NCC) reabsorbs 5-10% of filtered Na^+^ in the distal convoluted tubules (DCT). As highlighted by patients with Gitelman syndrome (loss-of-function mutations in NCC) and familial hyperkalemic hypertension (FHHt, also known as pseudohypoaldosteronism type II (PHAII)), NCC is essential for maintaining normal BP [4, 5]. NCC also plays an essential role in the antihypertensive effects of dietary K^+^, with the effects of a low K^+^ diet on BP absent in NCC knockout mice or mice receiving hydrochlorothiazide (NCC inhibitor) [6, 7].

Patients with Gitelman syndrome suffer from hypokalemia, while patients with FHHt suffer from hyperkalemia, demonstrating that NCC is also essential for K^+^ homeostasis [5, 8]. During hyperkalemia, NCC phosphorylation (active form) and abundance is reduced. This increases Na^+^ delivery to the amiloride-sensitive epithelial sodium channel (ENaC) [9] in downstream segments of the renal tubule, enhancing electrogenic K^+^ secretion via the renal outer medullary potassium channel (ROMK) and flow-dependent large Ca^2+^-activated K^+^ (BK) channels to help restore plasma K^+^ levels to normal. Conversely, during hypokalemia NCC phosphorylation and abundance are increased. Although this limits electrogenic K^+^ secretion, the consequence is increased Na^+^ reabsorption, hypervolemia and increased BP.

Studies in FHHt patients established that mutations in genes encoding for the With-no-lysine kinases 1 and 4 (WNK1 and WNK4), or the RING-type E3 ligase Cullin 3 (Cul3) and its substrate adapter Kelch-like protein 3 (KLHL3) control phosphorylation and activation of NCC [8, 10]. Mutations in WNK1 or WNK4 leads to hyper-activation of the STE20-related proline alanine rich kinase (SPAK) and Oxidative stress responsive 1 kinase (OSR1), with SPAK directly phosphorylating and activating NCC. Cul3 and KLHL3 together form an E3 ubiquitin ligase complex that ubiquitylates WNK1/4 kinases, targeting them for degradation. Loss-of-function mutations in Cul3 or KLHL3 prevent their interactions with WNK1/4, limiting degradation of these kinases and resulting in sustained activation of the WNK-SPAK/OSR1-NCC pathway.

Altered NCC phosphorylation following low or high dietary K^+^ intake is linked to alterations in the basolateral plasma membrane potential via the inwardly-rectifying potassium channel Kir4.1/Kir5.1 (a heterotetramer of Kir4.1 and Kir5.1 channels) and modulation of the WNK-SPAK/OSR1 kinase signaling pathway [11-15]. However, low K^+^ intake also increases phosphorylation of KLHL3, which impairs Cul3-WNK binding and degradation resulting in increased NCC [16]. Cul3 is one member of a large class of RING-type E3 ligases (Cul1-5, and Cul7 [17]). Activation of Cullins requires neddylation, a post-translational modification in which the ubiquitin-like modifier neuronal precursor cell-expressed developmentally down-regulated protein 8 (NEDD8) is covalently attached to the target protein [18, 19]. Processing of NEDD8 by the NEDD8-activating enzyme (NAE) is a prerequisite for binding of NEDD8 to Cullins [20, 21]. Conversely, enzymatic removal of NEDD8 from Cullins leads to deactivation of the ligase, a process regulated by the COP9 signalosome (CSN), an 8-subunit protein complex with isopeptidase activity. Specifically, the isopeptidase activity of CSN5 (or Jun activation domain-binding protein-1, Jab1) catalyzes the enzymatic removal of NEDD8 from the Cullin E3-ligase complex [22, 23].

Using mass spectrometry we recently discovered that Cul1 is enriched in the DCT, and Cul1 and Cul2 abundances are increased in the DCT of mice fed a high K^+^ diet for 4 days [24]. This suggests that cullins other than Cul3 may also play a role in mediating the inhibitory effects of K^+^ on NCC phosphorylation and abundance. To investigate this further, here we; 1) examined the abundance and neddylation status of cullins in mice fed different K^+^ diets or after incubation of *ex vivo* cultured renal tubules with K^+^; 2) examined NCC abundance and phosphorylation in Cul3-Δ9 mice (a model of FHHt with aberrant Cul3 function [25]) fed a high K^+^ diet; 3) examined in *ex vivo* isolated renal tubules the effects of high K^+^ on NCC phosphorylation and abundance during cullin or WNK kinase inhibition. Our data suggest that Cul3 and potentially other cullin family members are involved in the K^+^ mediated regulation of NCC.

## Experimental Procedures

### Animal experiments and tissue isolation

All protocols were approved and performed under a license issued for the use of experimental animals by the Danish Ministry of Justice (Dyreforsøgstilsynet). Male C57Bl/6J mice, 10-12 weeks of age, were kept in standard cages in a room with a 12:12-hrs artificial light-dark cycle with free access to tap water and a standard rodent chow (1324 pellets, Altromin, Germany). For experiments involving different K^+^ diets, mice were switched to either a control (1% K^+^) or high 5% K^+^ diet for 3 weeks, or a 0% K^+^ diet for 2 weeks (Teklad Diet, USA). Diets were prepared from powdered commercial diet free of Na^+^, K^+^ and Cl^-^ (Teklad Diet: TD.08251.Envigo. USA) and appropriate amounts of NaCl and KCl were added back to obtain 0% K^+^ diet (0.3% Na^+^, 0.3% Cl^-^ and 0% K^+^), 1% K^+^ diet (0.3% Na^+^, 0.3% Cl^-^ and 1.05% K^+^) and 5% K^+^ diet (0.3% Na^+^, 0.3% Cl^-^ and 5.25% K^+^). Mice were euthanized by cervical dislocation, followed by harvesting and homogenization of kidneys in ice-cold dissection buffer solution (pH 7.6) containing 250 mM sucrose, 10 mM triethanolamine, PhosSTOP, and cOmplete Mini tablets (Roche Diagnostics A/S, Mannheim, Germany). Kidney protein samples were prepared for immunoblotting using Laemmli sample buffer containing 15 mg/mL DTT.

### Studies in Cul3-Het/Δ9 mice

Cul3-Het/Δ9 mice, a model for FHHt, were generated as described [25]. Studies were approved by the Oregon Health and Science University Institutional Animal Care and Use Committee (protocol IP00286). Males aged 8-12 weeks were administered doxycycline (2 mg/ml in 5 % sucrose drinking water) for 2 weeks to induce Cul3-Het/Δ9 expression in epithelial cells throughout the renal tubule. Controls were genetically identical but received only 5 % sucrose drinking water. After a 2-week wash-out period mice were fed control diet (0.8% K^+^, TD.07309, Envigo) or a matched diet with 5% K^+^ (equimolar carbonate/citrate/Cl as anions; TD.07278, Envigo) for 7 days. The NaCl content was 0.32 % in both diets and water was provided ad libitum. Blood was collected via cardiac puncture under isoflurane anesthesia and transferred into heparinized tubes; 80 μl was loaded into a Chem8+ cartridge for electrolyte measurement by i-STAT analyzer (Abbot Point of Care Inc., Princeton, NJ, USA). Kidneys were harvested following blood collection under isoflurane anesthesia, snap frozen in liquid nitrogen, and stored at −80°C until homogenization. Sample processing and immunoblotting was performed as described [25].

### Kidney tubule suspensions

*Ex vivo* mouse kidney tubules preparations were generated as previously described [26]. Following isolation, tubules were equally divided into individual wells of tissue culture plates for further treatments. For experiments involving varying K^+^ concentrations, isolated tubules were initially incubated at 37 °C for 2 hours in media containing 3.5 mM K^+^ for acclimatization. After 2 hours, media was gently removed and replaced with new media containing either 2.5 mM, 3.5 mM or 6 mM K^+^ at 37 °C for 30 minutes or 24 hours. K^+^ was provided as KCl and NaCl was used to balance Cl^-^ concentration in the media. For experiments involving the general Cullin inhibitor MLN4924 (BioNordika) and varying K^+^ concentrations, media was gently removed and replaced with new media containing either vehicle (DMSO) or MLN4924 (0.5 μM) for 1 hour at 37 °C. Subsequently, media was removed and replaced with new media containing either 2.5 mM, 3.5 mM or 6 mM K^+^ with or without MLN4924 (0.5 μM) and tubules incubated at 37 °C for 30 minutes or 24 hours. For experiments involving the general WNK inhibitor stock2s (Tocris) and MLN4924, following acclimatization media was gently removed and replaced with new media containing 50 μM stock2s for 1 hour at 37 °C. Subsequently, media was removed and replaced with new media containing either vehicle or stock2s in the presence or absence of MLN4924 at 37 °C for 2 hours. Following all experiments, media was removed and protein was extracted from the tubules using Laemmli sample buffer containing 15 mg/mL DTT. Samples were sonicated and denatured at 60 °C for 15 minutes.

### Antibodies and immunoblotting

Primary antibodies used in immunoblotting include phosphorylated NCC (pT58) [27], NCC (SPC-402D, StressMarq), pSPAK/OSR1 (07-2273, Millipore), SPAK/OSR1 [28], Cul1 (sc-17775, Santa Cruz), Cul2 (sc-166506, Santa Cruz), Cul3 (sc-166110, Santa Cruz), Cul4 (sc-377188, Santa Cruz), Cul5 (sc-373822, Santa Cruz), CSN5/Jab1 (sc-13157, Santa Cruz), NAE (sc-390002, Santa Cruz), Proteasome 20S (ab3325, Abcam). Blotting of samples from Cul3-Het/Δ9 mice was performed using β-actin (ab8227, Abcam), NCC [29] and pT53NCC [29] antibodies. Specificity of the commercial antibodies was based on that they either gave a single unique band on an immunoblot corresponding to the target proteins predicted molecular weight, or the most prominent band on the immunoblot was at the target proteins predicted molecular weight (with no other bands of similar size). The pT58NCC, pT53NCC and total NCC antibodies have been validated using tissue from NCC knockout mice. Immunoblotting was performed as previously described [26], and for Cul3-Het/Δ9 mice [25]. Blots were developed using SuperSignal West Femto chemiluminescent substrate (Thermo Scientific, Denmark) or Amersham ECL Western Blotting Detection Reagent (GE Healthcare). Band intensity was quantified using Image Studio Lite (Qiagen) densitometry analysis.

### Statistics

Data in all graphs are shown as mean ± S.E.M. Individual sample size (n) is shown in figure legends. For comparing two groups of data, a Student’s unpaired t-test was used. For comparisons of more than two groups, one-way or two-way ANOVA followed by the Dunnett’s or Tukeys multiple comparison tests were used. Significance was considered at p < 0.05.

## Results

### High dietary K^+^ intake decreases NCC without major alterations in Cullin expression

Using mass-spectrometry, Cul1 and Cul2 expression were increased in the DCT of mice fed a high K^+^ diet for 4 days [24]. To further examine alterations in Cullin expression during high dietary K^+^ intake, mice were fed either a 0%, 1% or 5% K^+^ diet for 2-3 weeks (see *Methods*) and the abundance of Cullins in whole kidney homogenates determined using Western blotting. Relative to the 1% K^+^ intake, phosphorylation of NCC (pT58) and total NCC abundance were greater in the kidneys from mice receiving a 0% K^+^ diet, but lower in those receiving a 5% K^+^ diet (Fig. 1A and 1B). No significant differences in the abundances of Cul1, 2, 3, 4 and 5 were observed in whole kidney homogenates from mice under any of the dietary conditions (Fig. 1A and 1B). No significant differences were observed in the abundance of NAE (enzyme responsible for activation of Cullins), whereas CSN5/Jab1 levels (deactivates Cullins) were significantly higher on the 5% K^+^ diet relative to the 1% K^+^ diet (Fig. 1A and 1B).

**Figure 1.**
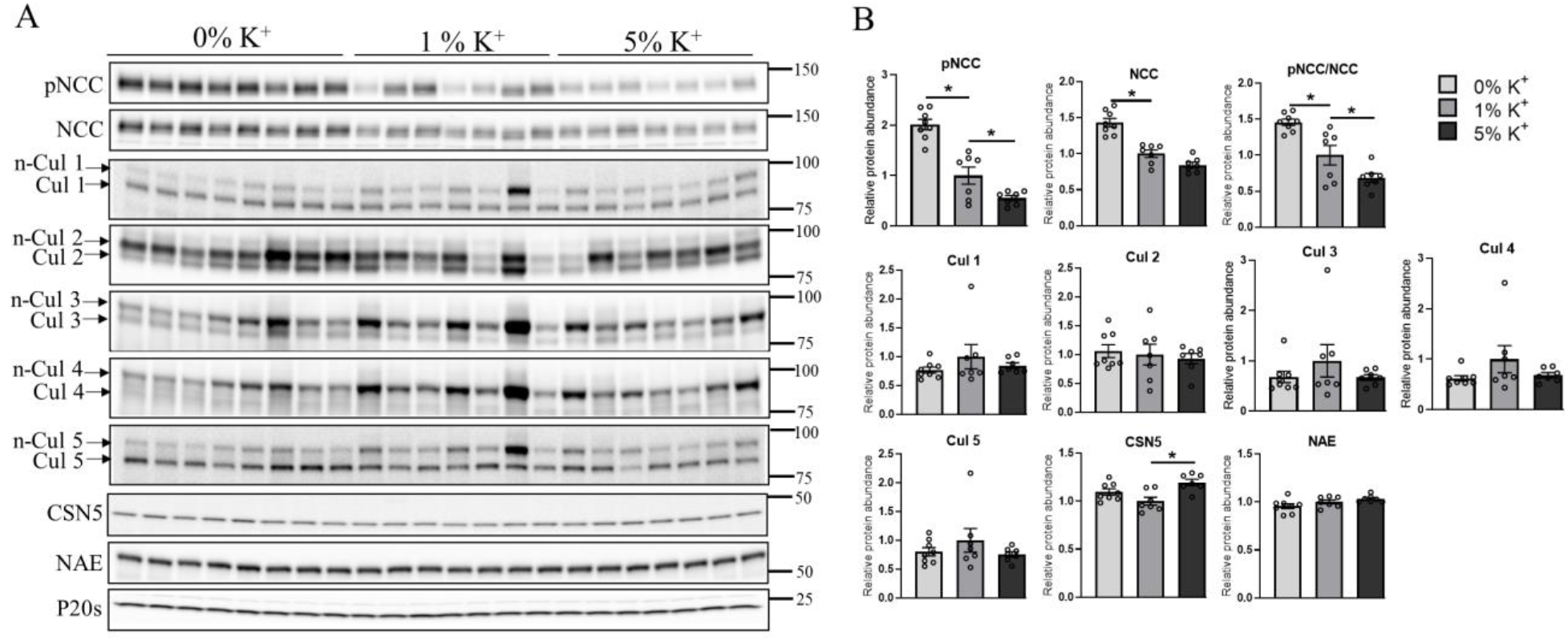
Increase in dietary K^+^ supplementation decreases renal abundance of phosphorylated and total NCC in mice without altering Cullin expression. A) Representative immunoblots of pNCC, NCC, Cullins (neddylated: n-Cul and non-neddylated: Cul), CSN5, NAE and P20s in kidney homogenates of mice placed in 0% K^+^ (n= 8), 1% K^+^ (n= 7) and 5% K^+^ (n= 7) diet. Molecular weight markers (kDa) are shown on right. B) Summarized relative protein abundance data. Each bar represents mean ± SEM. * indicates p <0.05 relative to mice that were on 1% K^+^ diet.

### Low dietary K^+^ intake is associated with decreased neddylation of Cullins

Cullin activity depends on their neddylation status, with cycling between neddylated and deneddylated forms changing the stability of the ligase, and the degree of neddylation having a direct correlation with their activity and ubiquitin-mediated degradation of their target substrates [30-35]. This neddylation status can be observed in the reducing conditions of Western blotting, where Cullin proteins are detected as a doublet (Fig. 1A); the lower molecular weight band representing the non-neddylated (inactive) form and the higher molecular weight band representing the neddylated (active) form. Here, the ratio between neddylated Cullins (n-Cul) to total Cullin abundance (Cul) was used to determine the percentage of active Cullins. In mice on a 0% K^+^ diet, neddylation levels of Cul1, Cul3, Cul4 and Cul5 were significantly lower than mice on a 1% diet (Fig. 2A). No significant changes to the neddylation status of Cul2 was observed following dietary manipulation (Fig. 2A). Non-linear regression analysis was applied to correlate the percentage change in pT58NCC and total NCC abundance following dietary K^+^ manipulation with the n-Cul/Cul ratio. A significant negative correlation was detected between pT58NCC or total NCC abundance and the n-Cul/Cul ratio for Cul1, 3, 4 and 5 (Fig. 2B and 2C). Although there appeared to be a positive correlation between the n-Cul2/Cul2 ratio and NCC, this did not reach significance (Fig. 2B and 2C).

**Figure 2.**
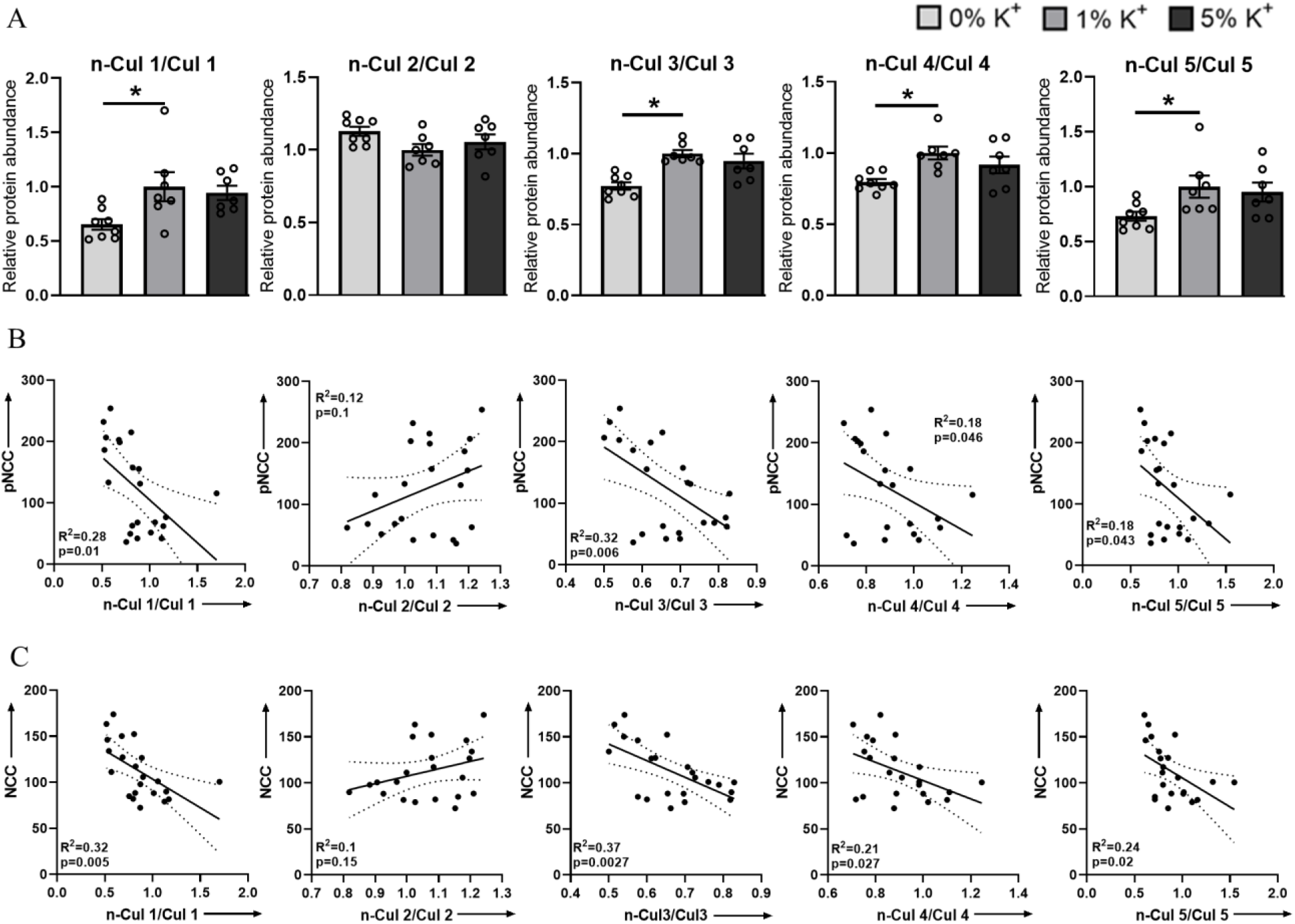
Low dietary K^+^ supplementation decreases neddylation levels of Cul1, 3, 4 and 5. A) Summary data of relative abundances of neddylated to non-neddylated form of Cul1, 2, 3 4 and 5. B) Linear regression analysis of n-Cul/Cul ratio and pNCC in kidneys of mice that were on different K+ diets (0% K^+^, 1% K^+^ and 5% K^+^). C) Linear regression analysis of n-Cul/Cul ratio and NCC in kidneys of mice that were on different K+ diets (0% K^+^, 1% K^+^ and 5% K^+^). Each bar represents mean ± SEM. * in panel A indicates p <0.05 relative to mice that were on 1% K^+^ diet.

### The effects of a high dietary K^+^ intake on NCC phosphorylation is attenuated in Cul3-Het/Δ9 mice

The Cul3-KLHL3 complex modulates NCC activity by altering ubiquitin-mediated degradation of WNK1 and WNK4 kinases [36]. Autosomal dominant mutations in the Cul3 gene cause haploinsufficiency and a severe form of FHHt, most likely a result of rapid Cul3 degradation following autoubiquitination [37]. This is recapitulated in a mouse model, CUL3-Het/Δ9 mice, which are characterized by higher levels of WNK kinases, and increased SPAK and NCC phosphorylation [10, 25, 37, 38]. To investigate if Cul3 plays a role in mediating the inhibitory effects of high K^+^ on NCC long-term, CUL3-Het/Δ9 mice were fed a control or high 5% K^+^ diet for 1 week. CUL3-Het/Δ9 mice had significantly increased renal expression of phosphorylated pT53NCC (100 ± 13.4% vs 288.5 ± 22.5%) and total NCC abundance (100 ± 11.8% vs 229.9 ± 30.8%) compared to the control animals (Fig. 3A and 3B). High dietary K^+^ significantly reduced pT53NCC (100 ± 13.4% vs 20.5 ± 3.1%; ∼ 80% reduction) and total NCC abundance (100 ± 11.8% vs 56.5 ± 3.2%; ∼ 44% reduction) in control mice (Fig. 3A and 3B). The high K^+^ diet significantly reduced total NCC abundance in CUL3-Het/Δ9 mice to a similar extent as observed in control animals (229.9 ± 30.8% vs 117.4 ± 8.9%; ∼ 49% reduction) (Fig. 3A and 3B). However, the reduction in pT53NCC expression in response to the high K^+^ diet in CUL3-Het/Δ9 mice, although significant, was not of the same magnitude as that observed in control mice (288.5 ± 22.5% vs 132.2 ± 23.5%; ∼ 54% reduction) (Fig. 3A and 3B). Furthermore, the phosphorylated NCC to total NCC ratio (as an indicator of relative NCC phosphorylation), was only significantly reduced in the control mice receiving a high K^+^ diet and not in CUL3-Het/Δ9 mice (Fig. 3A and 3B).

**Figure 3.**
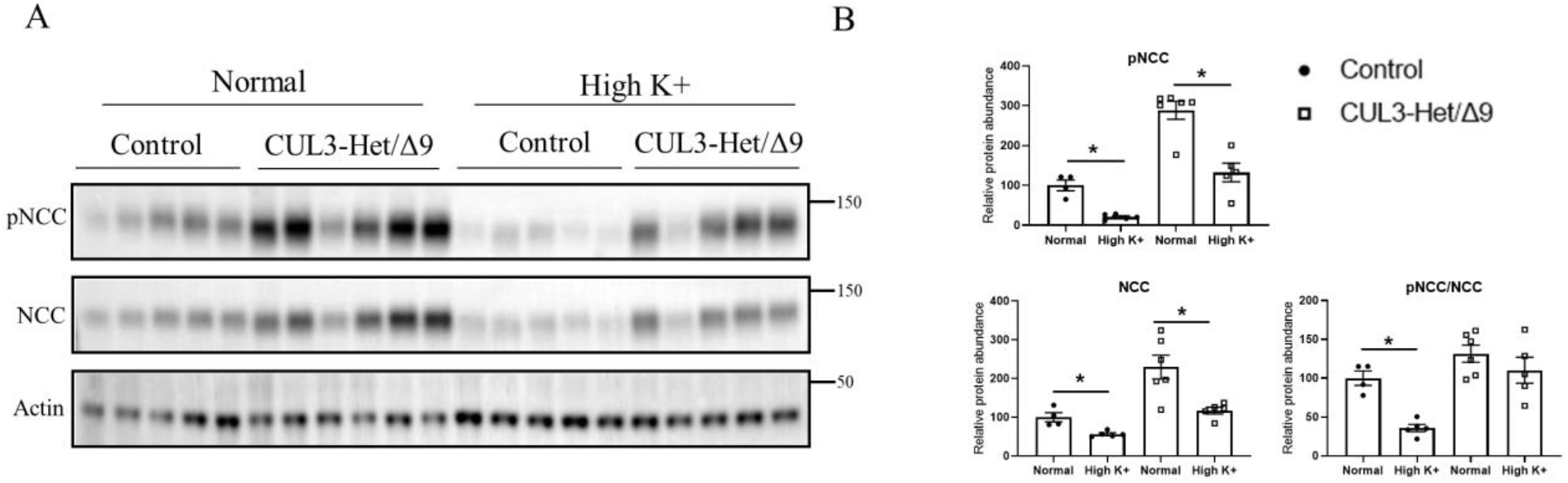
High dietary K^+^ supplementation effects on NCC phosphorylation is attenuated in Cul3-Δ9 mice. A) Representative immunoblots of pNCC, NCC and actin in kidneys homogenates of Control and Cul3-Δ9 mice that were on either normal diet (Control; n= 5 vs Cul3-Δ9; n= 6) or high K^+^ diet (Control; n= 5 vs Cul3-Δ9; n= 5). Molecular weight markers (kDa) are shown on right. B) Summarized relative protein abundance data. Each bar represents mean ± SEM. * indicates p <0.05 relative to mice that were on control diet in their respective group.

### Extracellular K^+^ changes NCC phosphorylation and abundance in isolated ex vivo renal tubules

Alterations in dietary K^+^ intake can alter levels of aldosterone, renin and angiotensin II, which likely play a role in modulation of NCC phosphorylation and abundance [39-41]. Altered plasma K^+^ levels *per se* observed during altered K^+^ intake also modulate NCC abundance and phosphorylation by modulation of the WNK-SPAK/OSR1 pathway [7]. To study the contribution of cullins in mediating K^+^ effects on NCC independent of other systemic factors, we used our extensively characterized *ex vivo* renal tubule preparations [24, 42, 43] and incubated them in different concentrations of K^+^ (2.5, 3.5 or 6 mM) for 30 min or 24 h (Fig. 4). After 30 min, compared to 3.5 mM K^+^, pT58NCC and phosphorylated SPAK levels were significantly increased and decreased after incubation in low or high K^+^ media, respectively (Fig. 4A and 4B). These changes occurred without significant differences in total protein abundances of NCC and SPAK (Fig. 4A and 4B). Similar effects on pT58NCC and phosphorylated SPAK were observed after 24 h incubation (Fig. 4C and 4D). After 24 h a significant difference in total NCC was also detected between tubules incubated in 2.5 mM and 6 mM K^+^. Despite these differences in NCC and SPAK levels, no significant differences were observed in the total abundance of Cul1, 3, 4 and 5 or in their n-Cul/Cul ratio at the time points examined (Suppl. Fig. 1).

**Figure 4.**
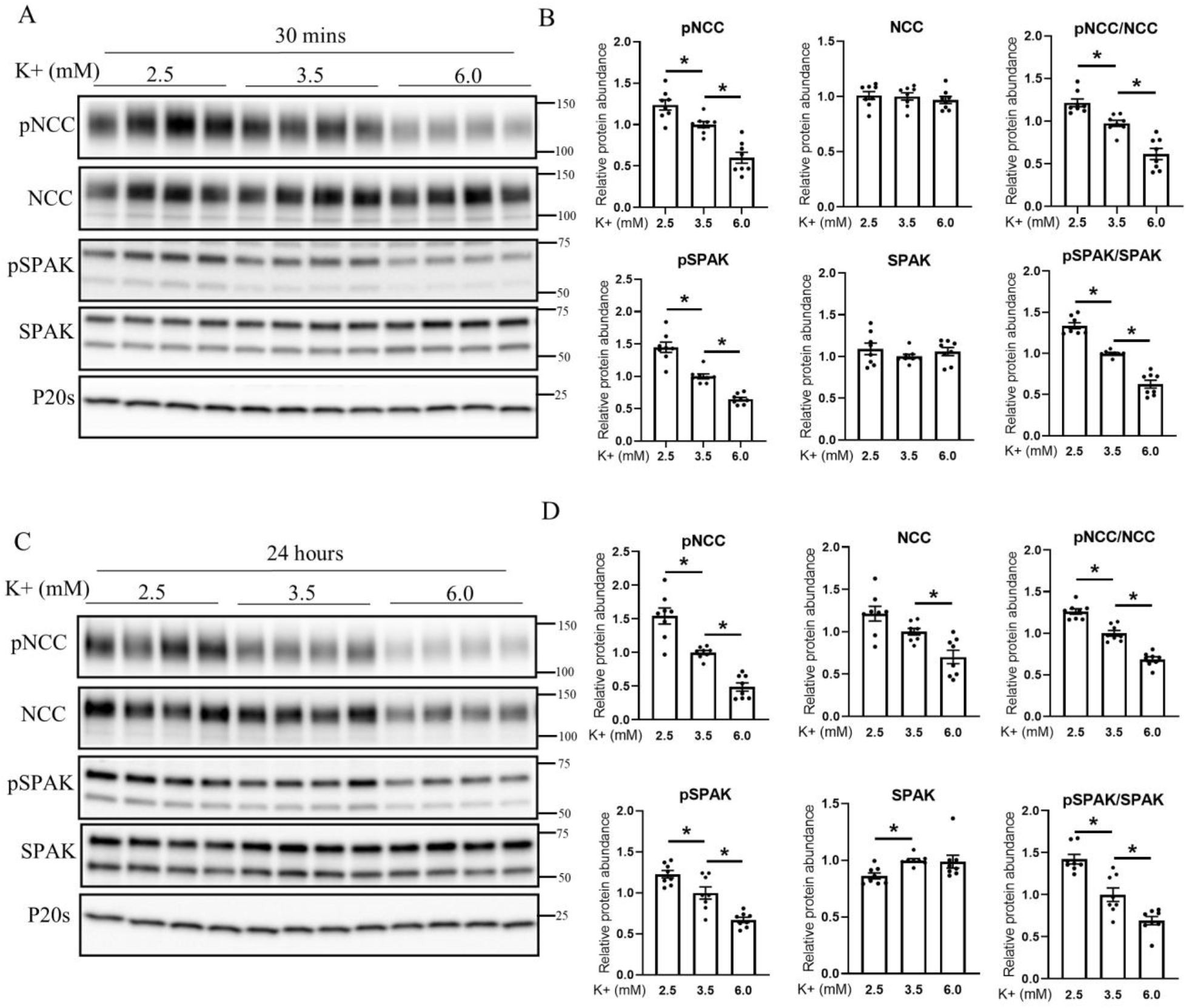
Extracellular K^+^ concentration regulates NCC phosphorylation in isolated renal tubules. A) Representative immunoblots of pNCC, NCC, pSPAK, SPAK and P20s in isolated renal tubules that were incubated in the media containing either 2.5mM, 3.5mM or 6mM K^+^ for 30 minutes *ex vivo*. Molecular weight markers (kDa) are shown on right. B) Summarized relative protein abundance data. C) Representative immunoblots of pNCC, NCC, pSPAK, SPAK and P20s in isolated renal tubules that were incubated in the media containing either 2.5mM, 3.5mM or 6mM K^+^ for 24 hours *ex vivo* and D) Summarized relative protein abundance data. Each bar represents mean ± SEM obtained from two independent experiments (n= 8). * indicates p <0.05 relative to 3.5mM condition.

### Pharmacological inhibition of Cullin activity ex vivo increases NCC phosphorylation

In *ex vivo* isolated tubules, the neddylation status (activity) of Cul1, 3, 4 and 5 under control conditions was different, suggesting they have different basal activities and capacities for activation or inhibition (Suppl. Fig. 2). However, treatment of tubules with MLN4924, a pan-Cullin inhibitor that prevents Cullin activation through inhibition of NAE [44], significantly decreased the neddylation state of Cul1, 3, 4 and 5 within 1 h, effects that remained stable over a 4 h period (Fig 5A and 5B). This occurred without changes in total cullin abundance (Suppl. Fig 3). Treatment with MLN4924 significantly increased SPAK and NCC phosphorylation compared to the vehicle treated groups, with phosphorylation levels increasing in a time-dependent manner (Fig. 5A and 5B). These changes in phosphorylation occurred without significant changes in total protein abundances.

**Figure 5.**
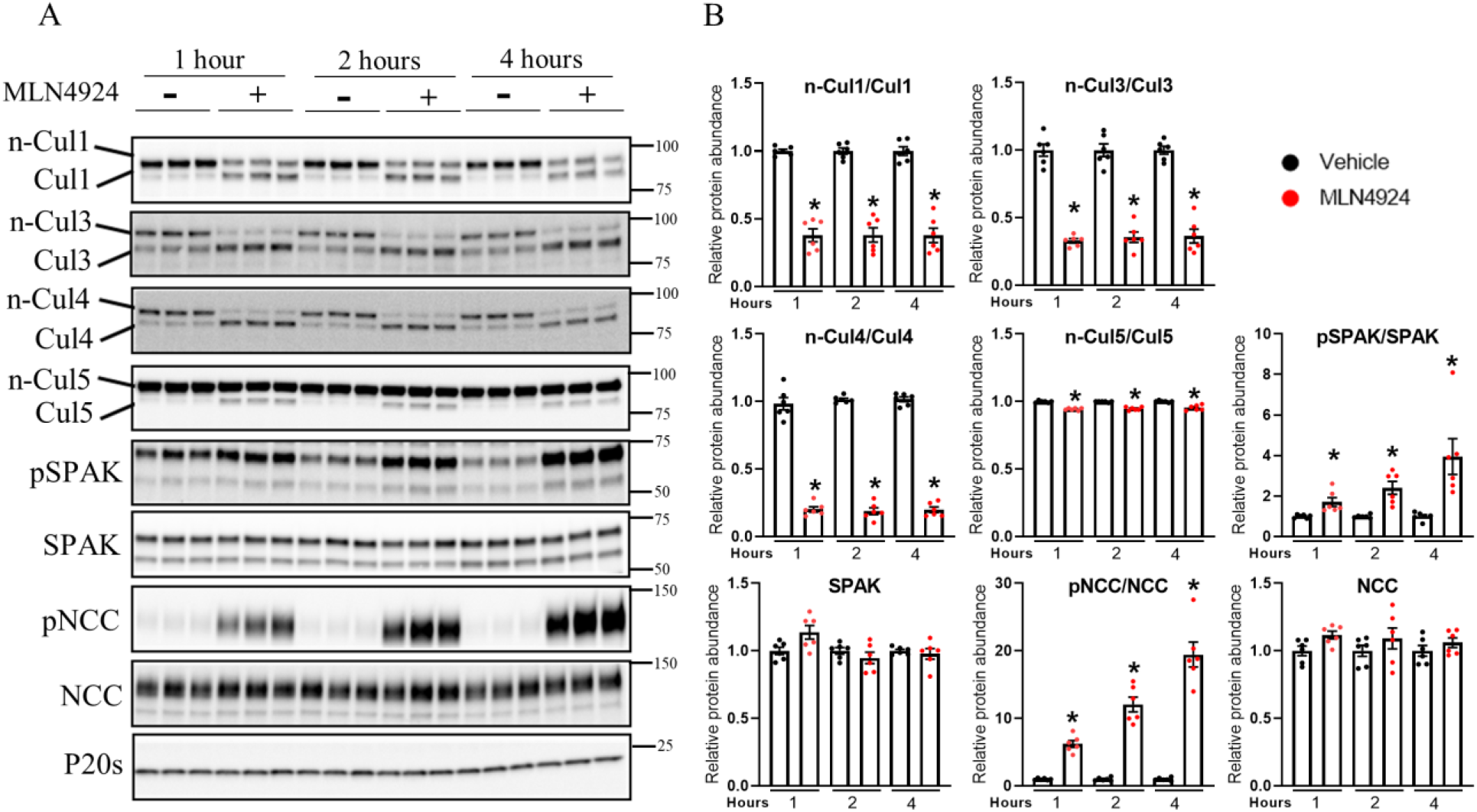
Pharmacological inhibition of Cullin neddylation increases NCC phosphorylation in isolated renal tubules *ex vivo*. A) Representative immunoblots of Cul1, Cul3, Cul4, Cul5, pSPAK, SPAK, pNCC, NCC, and P20s in isolated renal tubules that were incubated with either vehicle or Cullin inhibitor (MLN4924; 0.5uM) for various time points (1, 2 or 4 hours) *ex vivo*. Molecular weight markers (kDa) are shown on right. B) Summarized relative protein abundance data. Each bar represents mean ± SEM obtained from two independent experiments (n=6). * indicates p <0.05 relative to 3.5mM condition.

To further investigate if the effects of cullins on NCC are only via the alteration of the WNK-SPAK pathway, isolated *ex vivo* renal tubules were incubated with vehicle or the general WNK inhibitor (stock2s) for 1 h, followed by 2 h incubation in MLN4924 with/without WNK inhibition. As observed previously (Fig. 5), 2 h treatment with MLN4924 significantly increased SPAK and NCC phosphorylation compared to the vehicle treated groups (Fig. 6). However, in the presence of stock2s the effects of MLN4924 on SPAK and NCC phosphorylation were completely abolished (Fig. 6).

**Figure 6.**
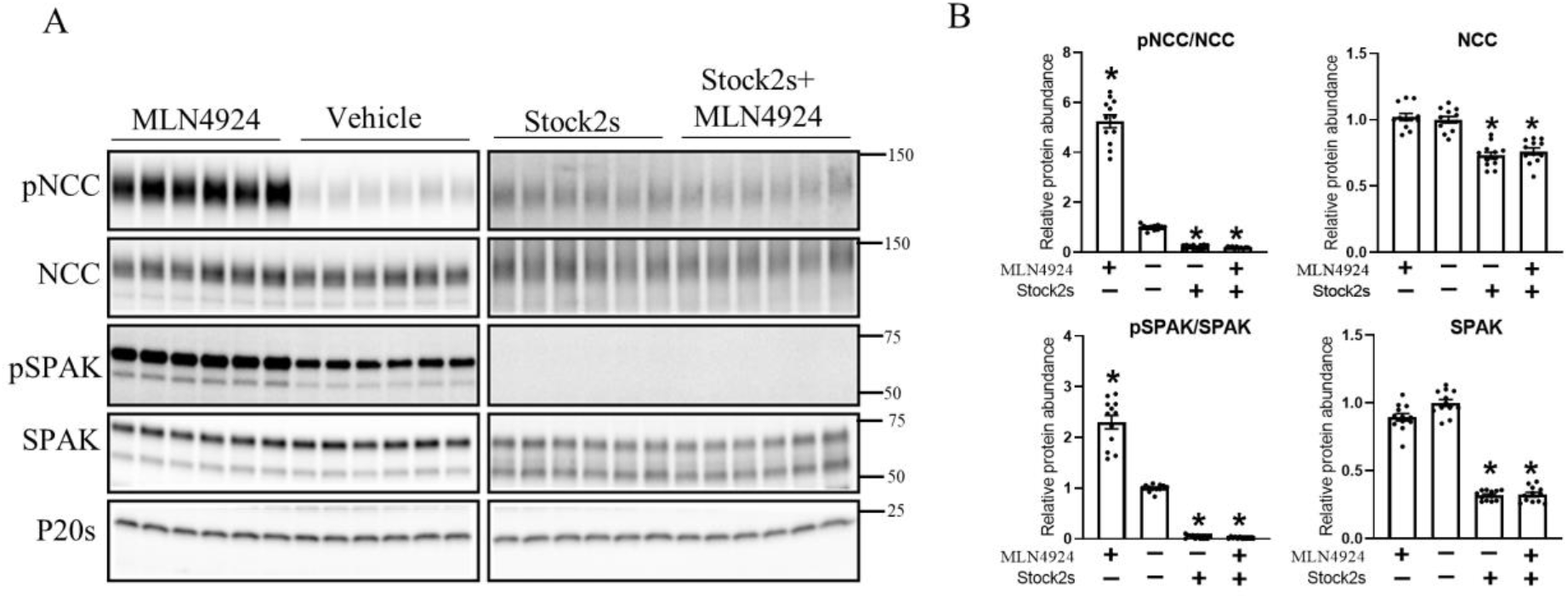
MLN4924 alters NCC phosphorylation via WNK-SPAK pathway. A) Representative immunoblots of pNCC, NCC, pSPAK, SPAK and P20s in isolated renal tubules that were incubated with either Cullin inhibitor (MLN4924), vehicle, WNK inhibitor (Stock2s) or both MLN4924 and WNK inhibitor. Molecular weight markers (kDa) are shown on right. B) Summarized relative protein abundance data. Each bar represents mean ± SEM obtained from two independent experiments (n= 12). * indicates p <0.05 relative to vehicle.

### The effects of K^+^ on NCC are attenuated during long-term cullin inhibition

To investigate the ability of K^+^ to modulate NCC in the absence of Cullin activity, isolated renal tubules were incubated in different concentrations of K^+^ (2.5, 3.5 or 6 mM) for 30 min or 24 h in the presence or absence of MLN4924. MLN4924 increased levels of NCC and SPAK phosphorylation without changes to the total protein abundance under all extracellular K^+^ conditions (Suppl. Fig. 4). As observed earlier (Fig. 4), when compared to tubules incubated in 3.5 mM K^+^, incubation with 2.5 mM K^+^ for 30 min significantly increased phosphorylation levels of SPAK and NCC, whereas levels were decreased in 6 mM K^+^ (Fig 7A and 7B). In the presence of MLN4924, phosphorylation levels of SPAK and NCC were not significantly increased after 30 min incubation in 2.5 mM relative to 3.5 mM K^+^, potentially because WNK and SPAK are already maximally active in the presence of MLN4924 (see *Discussion*). When tubules were incubated in 6 mM K^+^, SPAK and NCC phosphorylation were still significantly decreased (Fig 7A and 7B). No significant differences were observed in total protein abundances under any of the conditions. In contrast, after incubation of tubules for 24 h in 2.5 mM K^+^, the significant changes in total and pT58NCC levels (relative to 3.5 mM K^+^) were absent in the presence of MLN4924 (Fig. 8A and 8B). A significant reduction in total NCC was still observed when tubules were incubated in 6 mM K^+^ and MLN4924, but the change in phosphorylated NCC was attenuated, as emphasized by the lack of change in the phosphorylated NCC to total NCC ratio (like in CUL3-Het/Δ9 mice) (Fig 8A and 8B).

**Figure 7.**
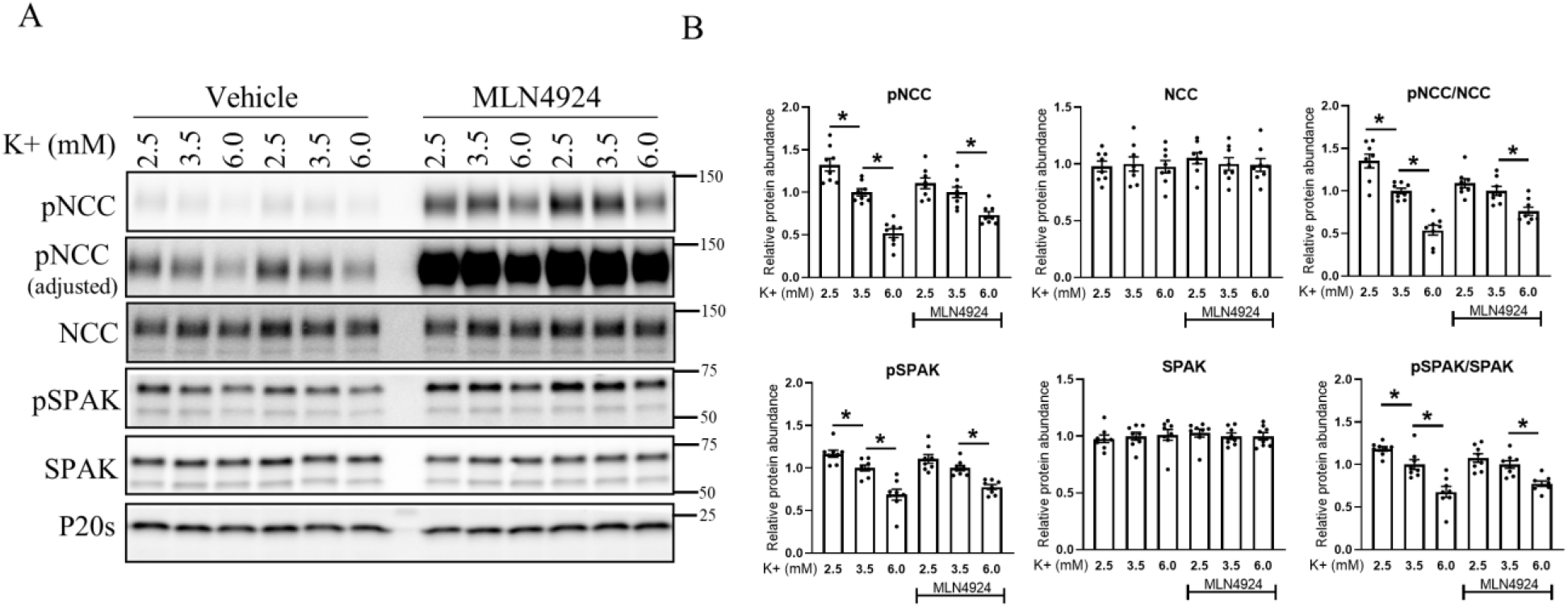
Increase in NCC phosphorylation in response to lower extracellular K^+^ is absent in the presence of MLN4924 in isolated renal tubules *ex vivo*. A) Representative immunoblots of pNCC, NCC, pSPAK, SPAK and P20s in isolated renal tubules that were incubated in the media containing different K^+^ concentration (2.5mM, 3.5mM or 6mM K^+^) in the presence of either vehicle or Cullin inhibitor (MLN4924; 0.5uM) for 30 minutes *ex vivo*. Molecular weight markers (kDa) are shown on right. B) Summarized relative protein abundance data. Each bar represents mean ± SEM obtained from two independent experiments (n= 8). * indicates p <0.05 relative to vehicle 3.5mM K^+^ group.

**Figure 8.**
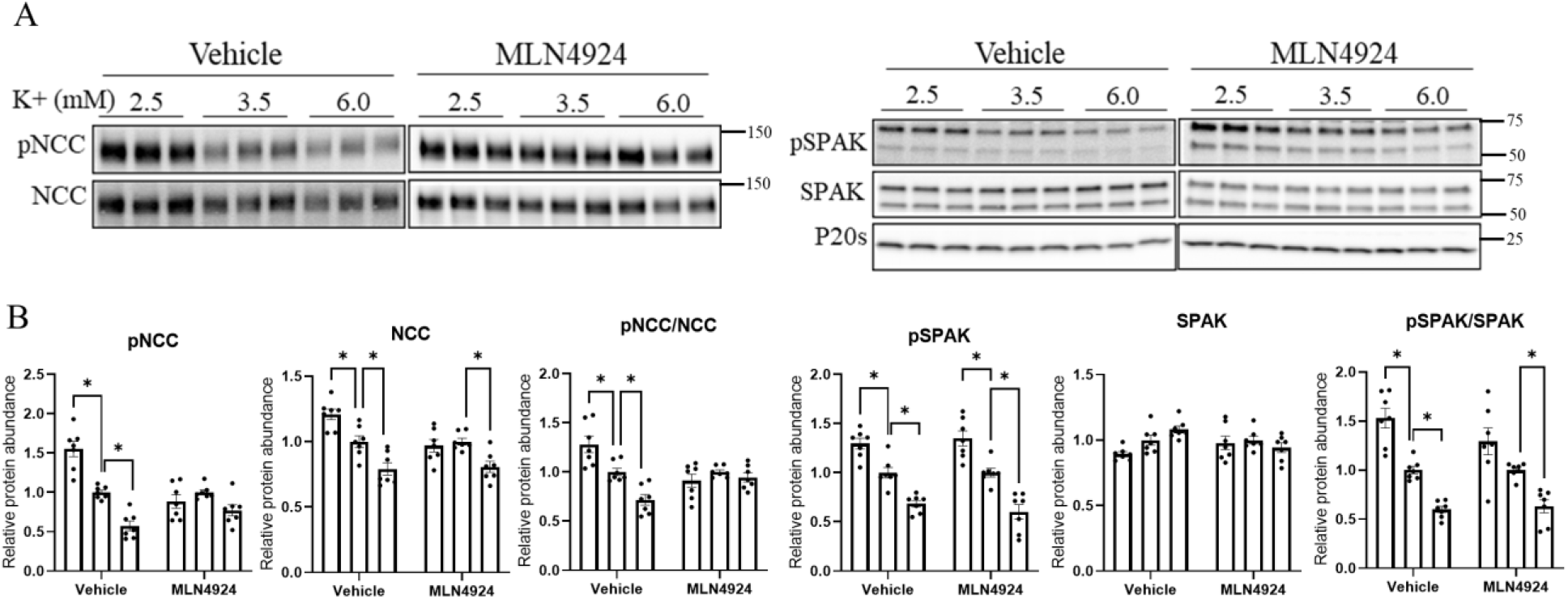
Lower extracellular K^+^ fails to induce NCC phosphorylation and abundance in the presence of MLN4924 in isolated renal tubules *ex vivo*. A) Representative immunoblots of pNCC, NCC, pSPAK, SPAK and P20s in isolated renal tubules that were incubated in the media containing different K^+^ concentration (2.5mM, 3.5mM or 6mM K^+^) in the presence of either vehicle or Cullin inhibitor (MLN4924; 0.5uM) for 24 hours *ex vivo*. Molecular weight markers (kDa) are shown on right. B) Summarized relative protein abundance data. Each bar represents mean ± SEM obtained from two independent experiments (n= 6). * indicates p <0.05 relative to 3.5mM K^+^ in their respective group.

We subsequently investigated if the absence of lower extracellular K^+^ effects on NCC expression in the presence of MLN4924 is due to altered expression of the basolateral heterodimeric K^+^ channel Kir 4.1/Kir 5.1. Relative to 3.5 mM K^+^, no changes in Kir 5.1 protein abundance were observed in isolated renal tubules incubated in 2.5 mM or 6 mM K^+^ for 24 h (Suppl. Fig. 5). Kir 4.1 protein abundance remained similar when tubules were incubated in 2.5mM K^+^ and 3.5 mM K^+^ (Suppl. Fig. 5), but it was significantly decreased in tubules incubated in 6 mM K^+^. However, this decrease in expression was also observed in the presence of MLN4924 (Suppl. Fig. 5).

## Discussion

Reduced dietary K^+^ intake is often associated with increased NCC activity and higher BP [2, 7, 45-48], whereas an increase in dietary K^+^ intake is usually associated with lower NCC activity and lower BP [2, 3, 49]. Dietary changes in K^+^ supplementation in mice alters WNK4 phosphorylation [7, 50] and WNK4^-/-^ mice do not increase NCC abundance after a low K^+^ diet [51], suggesting that K^+^ modulates NCC activity via alterations in the activity of WNK kinases. This alteration in WNK kinase activity is mediated, at least in part, through alterations in activity of the basolateral K^+^ channels Kir 4.1 and Kir 5.1 (forming a Kir4.1/5.1 heterodimer) and alterations in the intracellular concentration of Cl^−^ ([Cl^−^]_i_) [7, 13, 52, 53]. Supporting this, the inhibitory effects of a high K^+^ diet on NCC phosphorylation and abundance are greatly diminished in Kir4.1 or Kir5.1 knockout mice [14, 52] and the effects of acute K^+^ loading to reduce NCC phosphorylation are not observed in Cl^-^-insensitive WNK4 mice [54]. In addition to decreasing WNK4 phosphorylation, K^+^ supplementation reduces WNK4 abundance [7]. The Cul3 substrate adapter KLHL3 is important in this mechanism, with K^+^ dependent alterations in KLHL3 phosphorylation changing its ability to bind and degrade WNK4 [16, 55, 56].

Despite this body of evidence that alterations in K^+^ intake alter the activity and stability of WNK kinases [16], limited information existed on how K^+^ modulates the abundance or activity of cullins, and if inhibition of cullin activity alters K^+^-mediated effects on NCC. In an attempt to address this knowledge gap, here we investigated whether; 1) long-term alterations in dietary K^+^ intake change cullin abundance or their neddylation status, 2) short – or long-term K^+^ dependent changes in NCC abundance or phosphorylation independently of systemic hormones are dependent on cullin abundance or activity, and 3) if the effects of a long-term high dietary K^+^ intake on NCC are altered in a mouse model of FHHt mutation. Our results indicate that Cul3, and potentially other cullin family members, are important for modulating the overall response of the DCT to altered extracellular K^+^ and ultimately NCC activity (summarized in Fig. 9).

**Figure 9.**
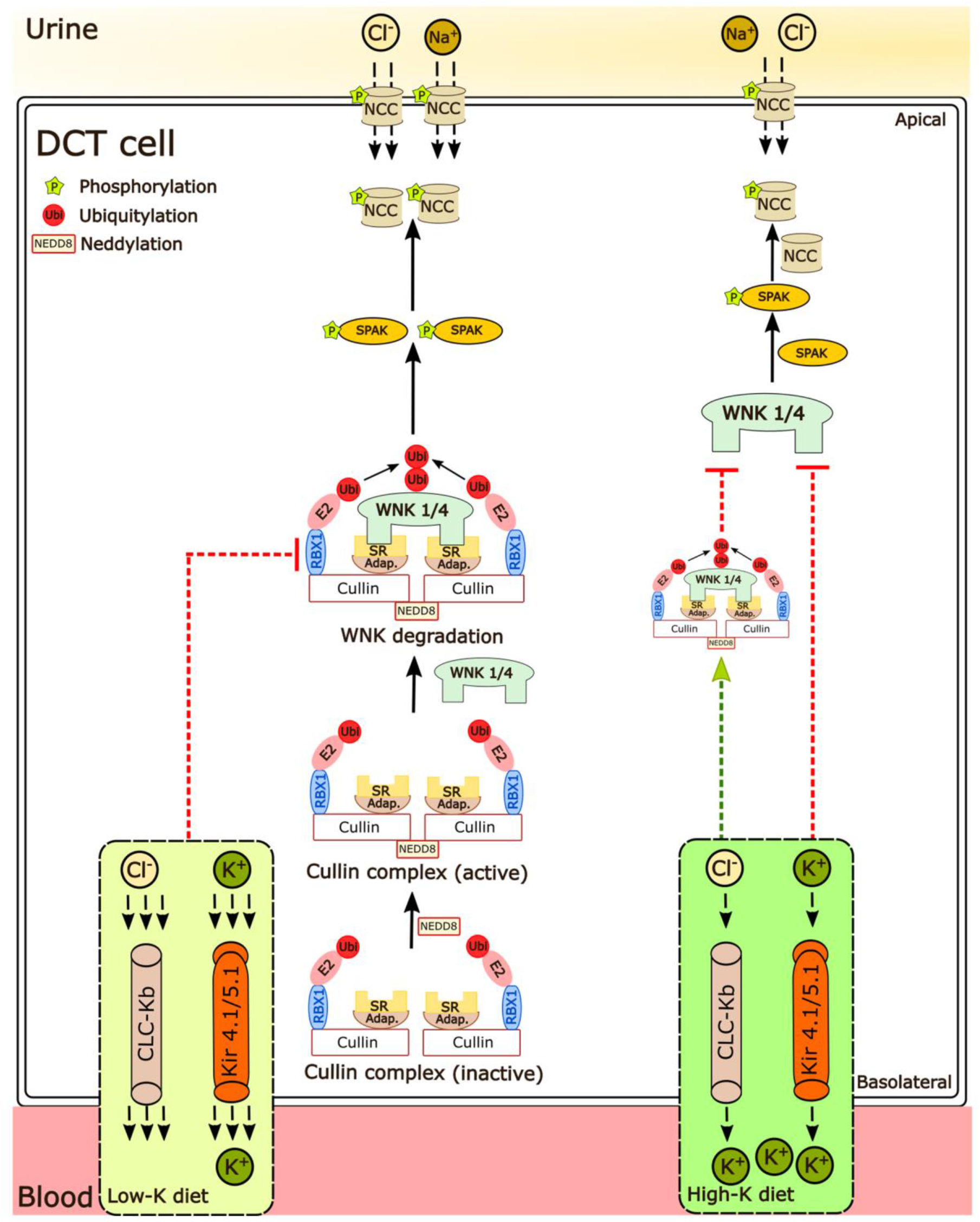
Regulation of NCC phosphorylation under different K^+^ conditions. Higher extracellular K^+^ concentration limits NCC phosphorylation, an effect partly dependent on cullin activity. Lower extracellular K^+^ concentration increases SPAK and NCC phosphorylation which is partly dependent on activity of cullins. SR-Substrate receptor; Adap.-adaptor protein.

In kidney samples from mice receiving a 0%, 1% or 5% K^+^ diet for 2-weeks, we observed progressively less NCC phosphorylation and abundance as dietary K^+^ intake increased (Fig 1.). These changes occurred without major differences in the total abundance of Cul1, 3, 4 and 5, and only a small change in the abundance of Cul2 (Fig. 1). In contrast, the neddylation status of Cul1, 3, 4 and 5 were increased as dietary K^+^ intake increased, which inversely correlated with phosphorylated and total NCC levels (Fig. 2). Together, this suggests that increased dietary K^+^ intake increases cullin activity, and this is associated with less NCC activity. As no significant changes in cullin neddylation were observed after incubating isolated renal tubules in different concentrations of K^+^ for 30 min or 24 h, systemic factors such as increased aldosterone may be mediating the altered cullin neddylation status observed *in vivo*, or longer periods of K^+^ exposure are required. A question that remains unanswered is how does the altered dietary intake alter cullin neddylation? We did not observe a difference in the abundance of the Nedd8 activating enzyme, but in principle altered abundance/activity of one or more components involved in the cullins neddylation/deneddylation pathway (CAND1, E1, E2 ligase and COP9 signalosome) or components of Cullin E3-ligase complex (E2 ligase and substrate receptors) could alter cullin activity. Interestingly, the Nedd8 E3 ligase DCN-like protein 4 (DCNL4) was recently identified as a K^+^ regulated protein modulating SPAK activity in kidney intercalated cells [57], but whether similar mechanisms occur in the DCT is unknown.

Inhibition of cullin neddylation *ex vivo* using MLN4924 potently increased SPAK and NCC phosphorylation within 1 h, an effect sustained over a 4 h period (Fig. 5). These changes were likely driven by increased WNK activity, as MLN4924 had no effect during WNK inhibition (Fig. 6). In the presence of MLN4924, no increases in NCC abundance (24 h) and phosphorylation (30 mins and 24 h) were observed in *ex vivo* tubules incubated in low K^+^ media (Fig. 7 and 8). Although this lack of response may suggest that K^+^-driven increases in NCC are dependent on cullin activity, the most likely explanation is that in the presence of MLN4924, SPAK and NCC phosphorylation are already at a maximum and no further increases are possible. These results are in line with what is observed in Cl^-^-insensitive WNK4 mice, where dietary K^+^ restriction did not further increase NCC phosphorylation above the already raised levels [54]. Together the results support that activation of the Kir4.1/5.1-WNK-SPAK pathway is the major mechanism for up-regulation of NCC by dietary K^+^ restriction [11-15]. However, as there is experimental evidence that cullins can interact and regulate various phosphatases, K^+^-mediated changes in protein phosphatase activity during dietary K^+^ restriction warrants further investigation [58-62].

After thirty minutes incubation of tubules in higher extracellular K^+^ (from 3.5 mM to 6 mM), NCC phosphorylation was still reduced despite the presence of MLN4924, suggesting acute reductions in NCC phosphorylation are independent of cullin and potentially WNK activity. This could suggest that in the short-term, the “on switch” for NCC and SPAK phosphorylation i.e. WNK kinase phosphorylation does not need to be “turned off” for the high K^+^ mediated effects to occur. However, acute effects of potassium on NCC phosphorylation are absent in Cl^-^-insensitive WNK4 mice and in SPAK knockout mice [54, 63], suggesting that *in vivo* there is a complex interplay between plasma K^+^, systemic factors and NCC that is absent in the *ex vivo* tubules. Such an idea is supported by acute K^+^ loading only decreasing NCC phosphorylation in mice that were prior K^+^-restricted for 1 day, but not in mice that were chronically K^+^-restricted for 5 days [63].

In contrast, in tubules the long-term effects of high K^+^ on NCC phosphorylation were attenuated in the presence of MLN4924, fitting well with studies in the FHHt CUL3-Het/Δ9 mice, where the ability of a high dietary K^+^ intake to reduce NCC phosphorylation was also limited (Fig. 3). Thus, when WNK4 activity is high, chronic effects of high K^+^ on NCC phosphorylation are reduced, but not prevented, suggesting that NCC dephosphorylation is also important. This is supported by studies in Cl^-^-insensitive WNK4 mice [54], where the chronic effects of potassium loading on NCC phosphorylation still occur. Together, our results and previous studies support that there are alternative time-dependent mechanisms at play on NCC during K^+^ loading, with early effects being driven by reductions in activity of the WNK-SPAK pathway, whereas later effects may be driven by alterations in the WNK-SPAK activity but also by changes in NCC dephosphorylation. Further studies are required to examine this possibility, but unfortunately the phosphatase responsible for NCC dephosphorylation subsequent to a high K^+^ load is still unclear with inconsistent results using PP1, PP2 or PP4 phosphatase inhibitors [46, 63-65].

## Supporting information

All supplemental

## Acknowledgements

We would like to thank the technical assistance of Tina Drejer.

## Authors’ contributions

RAF conceived the study. SKM, RL, SBP, JAM, DHE and RAF acquired data, analyzed and interpreted data. SKM and RAF drafted the manuscript. All authors’ critically revised manuscript for intellectual content. All authors approved the final version of the manuscript and agree to be accountable for all aspects of the work.

## Supplementary Material

Consists of **5** supplemental figures

