## Supplementary material for "Potassium effects on NCC are attenuated during inhibition of Cullin E3-ubiquitin ligases": All supplemental

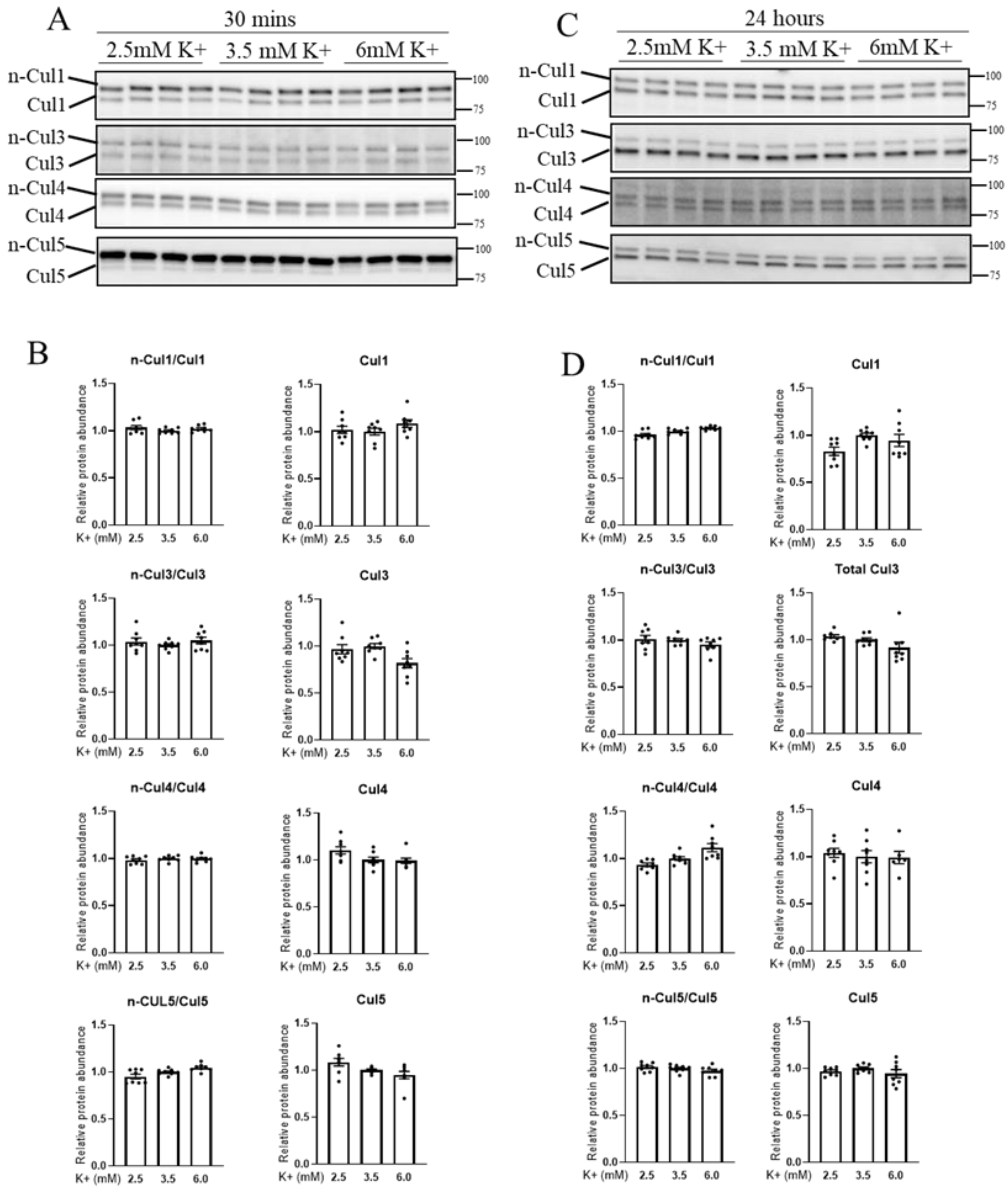

**Supplementary Figure 1. Incubation of ex vivo renal tubule preparations in media with various concentrations of $K^+$ has no significant effect on total abundance of Cul 1, 3, 4 and 5 or in their n-Cul/Cul ratio.** A and C) Representative immunoblots of Cul 1, Cul 3, Cul 4 and Cul 5 in isolated renal tubules that were incubated in media containing either 2.5-, 3.5-, or 6 mM $K^+$ for 30 min or 24 hours, respectively. B and D) Summarized relative protein abundance data from 30 min and 24 hours, respectively. Values are plotted as mean ± SEM with individual values shown (n= 8). For statistical analysis one-way ANOVA followed by the Dunnett's multiple comparison test was used. * indicates $p < 0.05$ relative to 3.5 mM condition.

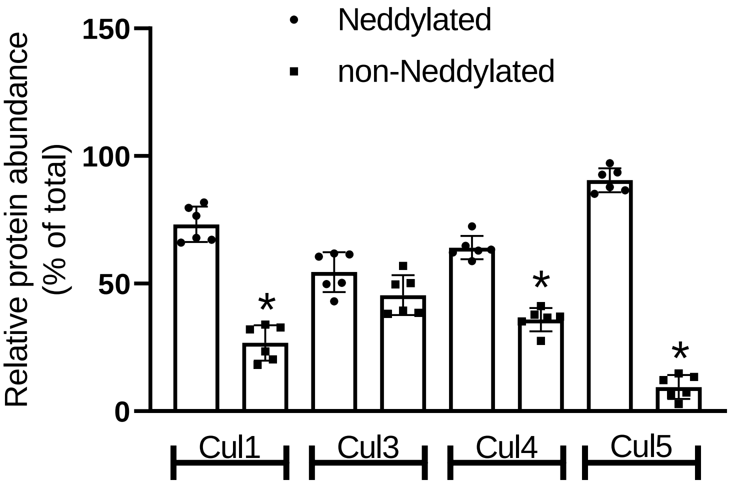

**Supplementary Figure 2. The basal neddylation status (activity) of Cul 1, 3, 4 and 5 are different**. The relative % of neddylated and non-neddylated Cul 1, Cul 3, Cul 4 and Cul 5 levels in isolated renal tubules are shown. * indicates $p<0.05$ relative to the neddylated group. For statistical analysis, a Student's unpaired t-test was used for individual comparisons of neddylated and non-neddylated Cullins.

A

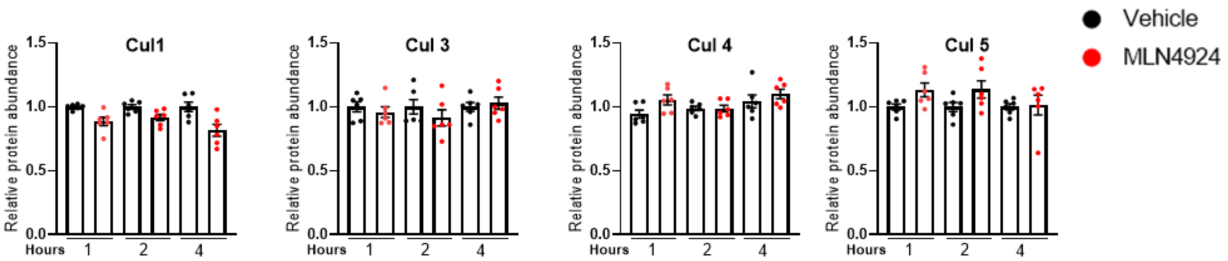

**Supplementary Figure 3. Treatment of isolated renal tubules ex vivo with MLN4924 had no significant effect on the total abundance of cullins**. A) Summarized data of Cul 1, Cul 3, Cul 4 and Cul 5 abundance in isolated renal tubules that were incubated with either vehicle or the Cullin inhibitor (MLN4924; 0.5 μM) for 1, 2 or 4 h. For statistical analysis, a Student's unpaired t-test was used for individual comparisons of vehicle and MLN4924 treated groups at individual time points.

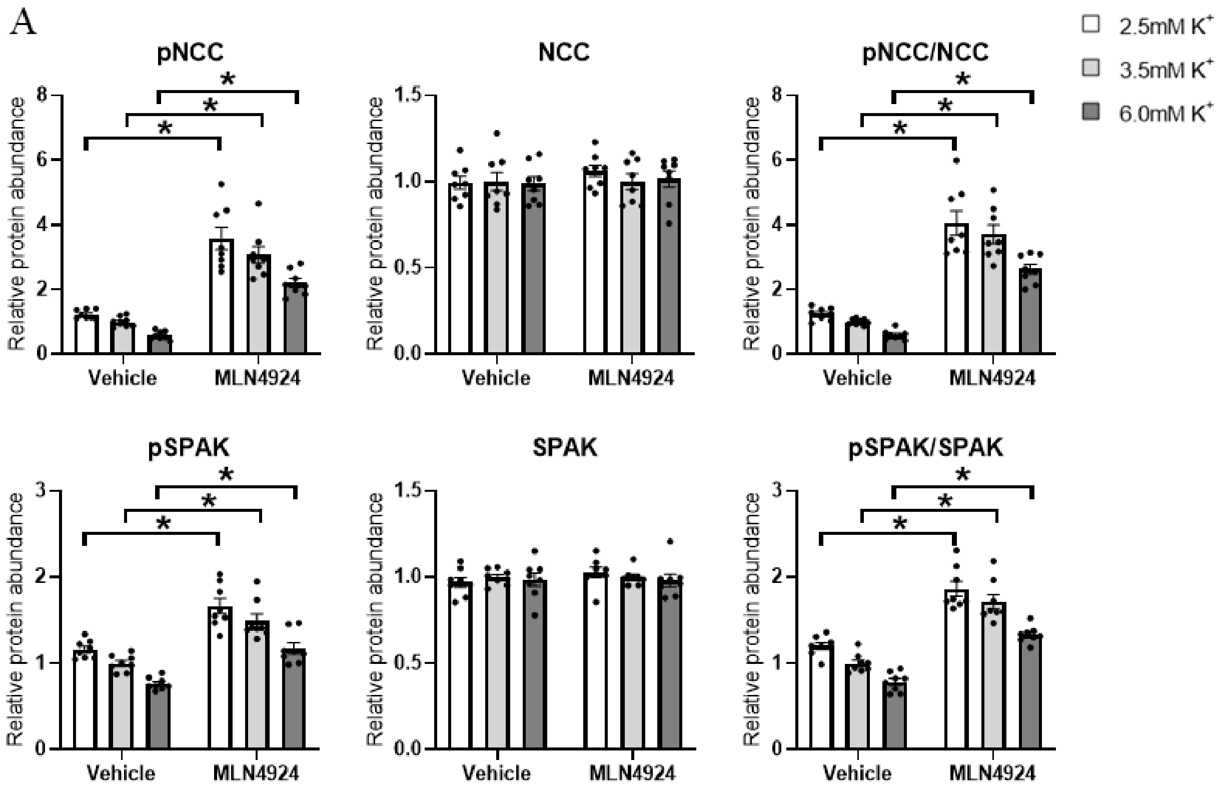

**Supplementary Figure 4. MLN4924 effects on NCC and SPAK phosphorylation under different extracellular K+ concentration.** A) Summarized data of pNCC, NCC, pNCC/NCC, pSPAK, SPAK and pSPAK/SPAK in isolated renal tubules that were incubated in media containing different $K^+$ concentrations (2.5 mM, 3.5 mM or 6 mM $K^+$) in the presence of either vehicle or the Cullin inhibitor (MLN4924; 0.5 μM) for 30 minutes. All data are normalized to vehicle 3.5mM $K^+$ group. * indicates $p<0.05$ relative to vehicle 3.5mM $K^+$ group. For statistical analysis Two-way ANOVA followed by the Tukey multiple comparison test was used.

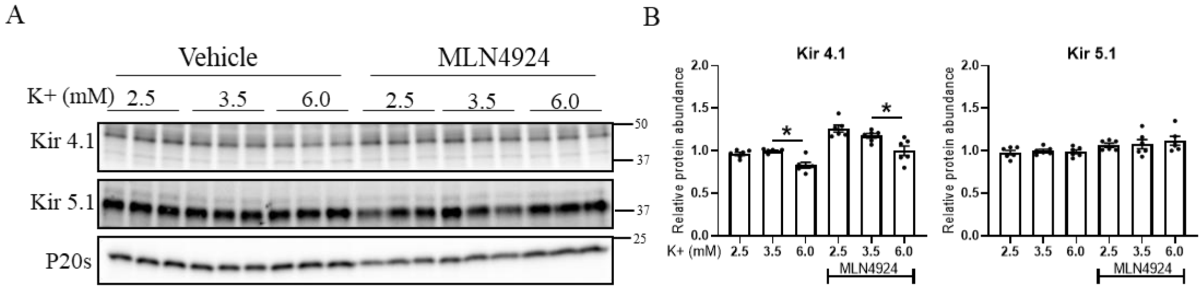

**Supplementary Figure 5. Inhibition of cullins does not alter $K^+$ effects on the $K^+$ channels Kir 4.1 and Kir 5.1.** A) Representative immunoblots and B) summarized data of Kir 4.1 and Kir 5.1 abundance in isolated *ex vivo* renal tubules that were incubated in media containing different $K^+$ concentrations (2.5 mM, 3.5 mM or 6 mM $K^+$) in the presence of either vehicle or the Cullin inhibitor (MLN4924; 0.5 μM) for 24 hours. * indicates $p<0.05$ relative to 3.5 mM $K^+$ in their respective group. For statistical analysis one-way ANOVA followed by the Dunnett's multiple comparison test was used.
